# Social dominance status and social stability in spiny mice (*Acomys cahirinus*) and its relation to ear-hole regeneration and glucocorticoids

**DOI:** 10.1101/2022.09.13.507818

**Authors:** Justin A. Varholick, Gizelle Godinez, Sarim Mobin, Ashley Jenkins, Russell D. Romeo, Jacob Corll, W. Brad Barbazuk, Malcolm Maden

## Abstract

Spiny mice (*Acomys cahirinus*) are an emerging animal model in studies measuring tissue regeneration, but decades of research on social dominance in other animals indicates the relationships animals form in their home-cage may affect phenotypic plasticity in tissue regeneration and glucocorticoids. Studies in baboons and mice, for example, indicate that subordinate ranked animals heal wounds slower than their dominant group-mates, and have increased levels of basal glucocorticoids. Recent studies in tissue regeneration with salamanders and zebrafish indicate that increased glucocorticoids can delay tissue regeneration, but whether this effect extends to *Acomys* is unknown, especially regarding their social dominance relationships. Here we report that most adult *Acomys* had a social dominance status, but many groups had unclear social stability, with more frequent huddling than fighting during their active cycle. We also found no sex differences in social dominance behavior, and that *Acomys* more frequently fled than froze when chased or approached. After a 4mm ear-pinna biopsy, we found that social stability significantly accounted for variability in time to close the ear-hole but adding age to the statistical model removed the effect of social stability. When investigating glucocorticoid blood levels, there were no significant effects of social dominance status or social stability. A transcriptional enhancer for StAR, Nr5a1 had a significant effect for the interaction of social dominance status and social stability. This effect, however, was not reflected in StAR and unclear groups mostly had unclear social statuses, so this effect should be considered with caution. This is the first study to investigate home-cage social dominance behaviors in *Acomys* since the 1970s or measure any associations with their ability to regenerate tissue. This provides a platform for further work on their social dominance and glucocorticoids and highlights the need to consider the role of aging in their ability to regenerate tissue.

## Introduction

Social relationships often form in groups of animals as they engage in agonistic behaviors while competing for resources in their environment (e.g., space/territory, food, water, or mates) (1,2). Over time, a predictable dominance relationship can form where one animal consistently yields—the subordinate—while their partner historically attacks, injures, or gains priority access to the resource—the dominant (3). An individual’s social dominance status, however, can be unstable, with animals switching rank, or be unclear with animals frequently fighting or never fighting (4,5). Engaging in agonistic behavior directly activates the sympathetic nervous system to release epinephrine and norepinephrine, and the hypothalamic-pituitary-adrenal (HPA) axis to release glucocorticoids (6). Repeated or chronic activation of the HPA axis can be particularly damaging and is associated with metabolic syndrome, obesity, cancer, cardiovascular disease, and susceptibility to infection (6). Indeed, subordinate animals—that more frequently yield—can have higher levels of basal glucocorticoids (e.g., cortisol or corticosterone) in their blood (7,8), higher levels of inflammatory cytokines (9), and delayed wound-healing (10,11). Subordinate animals, however, are not the only ones susceptible to these effects. Dominant animals, for example, can also experience chronic stress if they are frequently fighting to maintain or reinforce their position, and all animals may experience chronic stress when living in unstable social groups (12). Thus, phenotypic diversity in glucocorticoids, inflammation, and wound healing can arise within and between social groups depending on social dominance status and social stability.

Although it is well recognized that social relationships are a central aspect of life and coincide with phenotypic diversity, these relationships are often neglected from experimental designs and statistical plans (4,13–15). In general, scientists attempt to standardize—or reduce the diversity of—any hereditary, environmental, or developmental factors except their treatment. While this practice not only neglects the inherent phenotypic diversity fundamental to biology, it is unrealistic and contributes to spurious findings or irreproducible results (16–19). For example, in the interest of standardization, scientists may assume that all animals cage are exposed to the same environment (14). With regards to standardization it would be more ideal to house the animals alone to remove any chance that the cage-mates influence phenotypic traits in the experiment, but we recognize that solitary housing greatly departs from their phylogenetic and developmental history, and thus may give biased and non-generalizable results (20). We therefore house animals in social groups to account for their social needs, and ignore the effect of individual cage-mates on phenotypic traits to account for our standardization needs (4,21). Recent studies indicate, however, that social dominance status and social stability can more consistently account for phenotypic diversity than the physical cage-context (4), and whether social context interacts with an experimental outcome greatly varies between experiments (22). Thus, it is imperative that we begin understanding the social contexts of our lab animals and how it may interact with our treatments and phenotypic traits of interest.

Spiny mice (*Acomys cahirinus*) are social animals known to form social dominance relationships (23) and are an emerging model system in the field of tissue regeneration (23–26)—a field measuring how animals heal after injury. Over the past decade, scientists have determined that spiny mice can regenerate a number of tissues and organ systems in response to injury (e.g., removal or transection of skin, hair follicles, cartilage, muscle, nerve, and spinal cord) (24,27). Many studies using *Acomys* injure their ear pinna with a 4mm biopsy punch and find that the skin, hair follicles, fat, and cartilage regenerate without scarring, while lab mice show little to no regeneration and scar (28–31). This is an exciting discovery because mammals were previously thought to have impaired regenerative abilities, often healing tissues with a scar and reduced functionality rather than the remarkable ability of salamanders to regrow appendages and zebrafish to regenerate heart tissue (32,33). While this ability to regenerate is remarkable compared to the healing abilities of common mammals, recent evidence indicates that increased glucocorticoids are associated with delayed or disrupted tissue regeneration in salamanders and zebrafish (34–36). While the direct mechanisms underlying the relationship between increased glucocorticoids and delayed or disrupted regeneration remains unclear, it is also unknown whether this phenomenon applies to other regeneration-competent animals like *Acomys*.

The role of glucocorticoids on tissue regeneration in *Acomys* is particularly important because they are known to form social dominance relationships and these relationships in other animals are associated with phenotypic diversity in glucocorticoid levels and healing (10,11,37). Thus, any diversity in glucocorticoids due to social dominance relationships could also be associated with diversity in regeneration. Currently our understanding of the social dominance relationships of *Acomys* are limited to a few studies. The initial studies on their agonistic behavior were published in the late 1970s and had two major findings: i) both males and females engage in overt forms of agonistic behavior like chasing and attacking, and ii) the females are often dominant in mixed-sex housing (38,39). Overt forms agonistic behavior are usually limited to male rodents (2), with females showing more covert forms like side-pushing or over-climbing (40). However, female *Acomys* are not the only exclusion to this rule. Other female rodents like the golden hamster (*Mesocricetus auratus*) also more typically engage in overt forms of agonistic behavior and are dominant in mixed-sex housing (41–43). Nonetheless, most studies on the relationship between glucocorticoids and social dominance are limited to common rodents where males are more likely to engage in overt forms of agonistic behavior and females more likely to engage in covert forms. Thus, previous studies in common rodents showing a relationship between social dominance and glucocorticoids may not reflect the same relationship in *Acomys*, and thus warrant investigation, especially given the role of glucocorticoids on regeneration in other species.

The original study on social dominance behavior in *Acomys* also noted that they show a general lack of freezing behavior or “appeasement gesture(s)” (38). This behavior is also unusual for a rodent (38), as they are thought to more commonly freeze or show a subordinate posture like lying on their back when repeatedly attacked or chased (2,44,45). Freezing and subordinate postures often correspond to a de-escalation of agonistic behavior, lower the risk of injury, and likely activate the HPA axis differently than continuing to chase and flee. Thus, *Acomys* are either not de-escalating agonistic behavior over time or have alternative and more cryptic forms of subordinate agonistic behavior, which will have a differential effect on HPA axis activation and glucocorticoid regulation. Notably, some studies on *Acomys* prey behavior also find that they show a general lack of freezing during a predatory attack (46–48), and no significant behavioral response to an predatory call (49). When investigating for differences in glucocorticoids during these situations, however, they find that although *Acomys* show no significant behavioral response to a predatory call, they do have significantly increased glucocorticoid levels compared to baseline and a human voice (49). Thus, *Acomys* are responding to the predator with an expected glucocorticoid response and an unexpected behavioral response.

Glucocorticoids are also demonstrably different in *Acomys* compared to more common rodents like mice and rats. First, cortisol is the primary glucocorticoid in *Acomys*, whereas corticosterone is the primary glucocorticoid in mice and rats (50,51). Female *Acomys* also tend to have higher levels of glucocorticoids (52,53), but these effects are unclear in the literature (50) and could be due to differences between cage-groups (53). It remains unclear if *Acomys* with different social dominance statuses differ in glucocorticoid levels, like other animals, and whether differences between cage-groups is due to differences in social structure or stability.

The current study investigated social dominance in *Acomys* to determine whether differences in social dominance are also associated with differences in healing and glucocorticoids, as observed in other mammalian species, to explore whether their social contexts should be considered in experimental designs and statistical plans. We hypothesized that all same-sex and established groups of adult *Acomys* (in dyads or triads) would have stable social dominance statuses, with no significant differences in agonistic behaviors between sexes, ages, or group-sizes, and they would rarely freeze during agonistic interactions. We also hypothesized that subordinate *Acomys* would have delayed ear-hole regeneration and increased glucocorticoid hormones (i.e., cortisol measured by radioimmunoassay) along with increased gene expression of genes involved in the synthesis of glucocorticoids (i.e., Cyp11a1, Cyp11b1, and StAR measured by real-time quantitative polymerase chain reaction (RTqPCR)). We also explored differences in Nr0b1 and NR5a1, transcriptional enhancers of StAR, with the prediction that they would also be increased in subordinate animals.

## Results

### Social dominance status and asymmetry of dominance relationships

First, the social dominance status and asymmetry of the dominance relationships were determined across nine days, three per week for three consecutive weeks. The groups available for this study were housed in 4 different housing conditions: young female dyads (YFD), young male dyads (YMD), young male triads (YMT), and old female dyads (OFD)—three social groups per housing condition. Dominance status was determined by calculating a David’s Score, which measures the individual proportion of wins, or offensive agonistic behaviors, across the observation time. Positive David’s Scores represent a higher proportion of wins, while negative scores represent a higher proportion of losses, or defensive agonistic behaviors. In most cases, cage-mates could be assigned a unique social dominance status of dominant, subdominant, or subordinate—depending on group-size (Fig. 1A). However, some cages had very similar proportions of winning dominance interactions (measured as David’s scores) with high asymmetry ≥ 0.75 in the directionality of the dominance interactions (i.e., directional consistency (DC) index) (i.e., Cages B, H, and I). Older females in dyads did not engage in any agonistic behavior during our video-recordings and were thus assigned an unclear dominance rank. Comparison of DC indices determined there were no significant differences between the sexes (*W*(3,3)=8, p=0.200) or group-sizes for males (*W*(3,3)=9, p=0.350) (Fig. 1B).

**Fig. 1:**
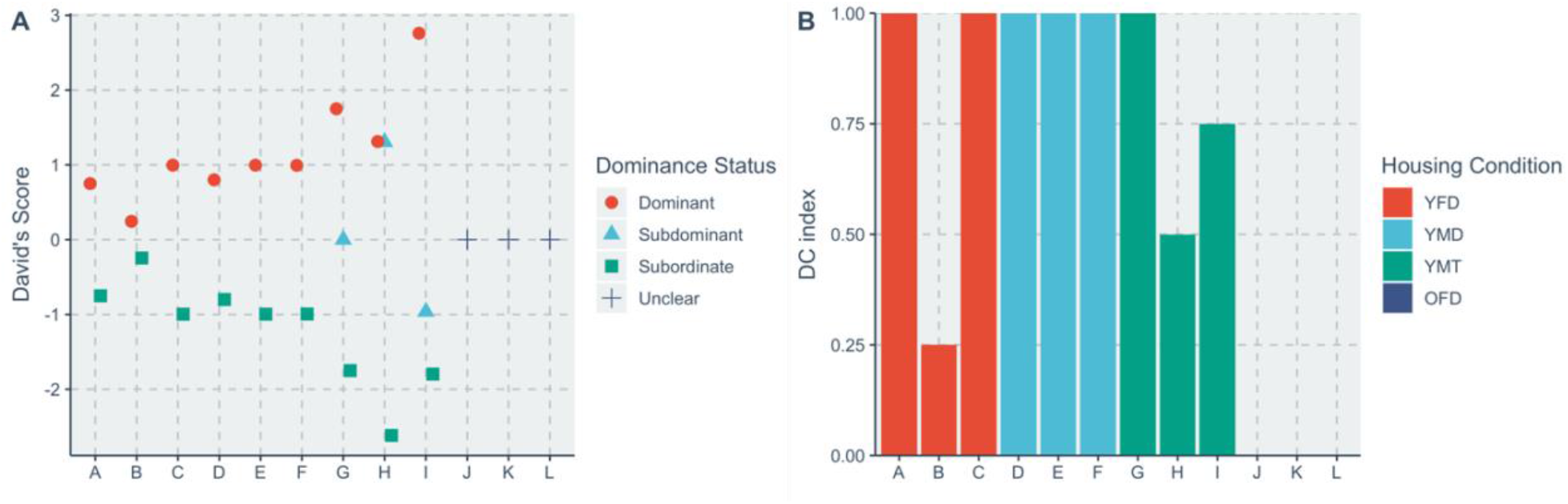
David’s scores and DC indices across cages. **A**: Scatterplot of individual David’s scores across all three weeks for each cage labeled with a letter. **B**: A bar graph of each cage’s DC index, groups colored by housing condition.

### Stability of social dominance status

Weekly David’s scores and social dominance status assignments were determined by splitting the agonistic behavior by week, to determine group stability. Unstable groups were defined as any groups that changed social dominance status at any point during the three-week observation. Those that maintained the same social dominance status across weeks were defined as stable. Notably, groups that had a defined social dominance status that later became unclear in subsequent weeks because of no fighting were also determined as unclear rather than unstable. Six groups had an unclear dominance status across weeks (i.e., Cages A, D, G, J, K, and L), while three groups were stable (i.e., Cages C, E, and F), and three groups were unstable (i.e., Cages B, H, and I) (Fig. 2). Notably, only the dominant *Acomys* in the unstable Cage of “I” remained stable throughout the 3 weeks. The instability observed across weeks was also supported by the cages’ closer David’s scores and DC indices ≤ 0.75 (see previous section). An exploratory analysis found that age was a significant predictor of social stability (*F*(1,10)=7.146, p=0.023) with older *Acomys* having an increased chance of being unclear, in a linear mixed effects model with cage-identity as a random factor.

**Fig. 2:**
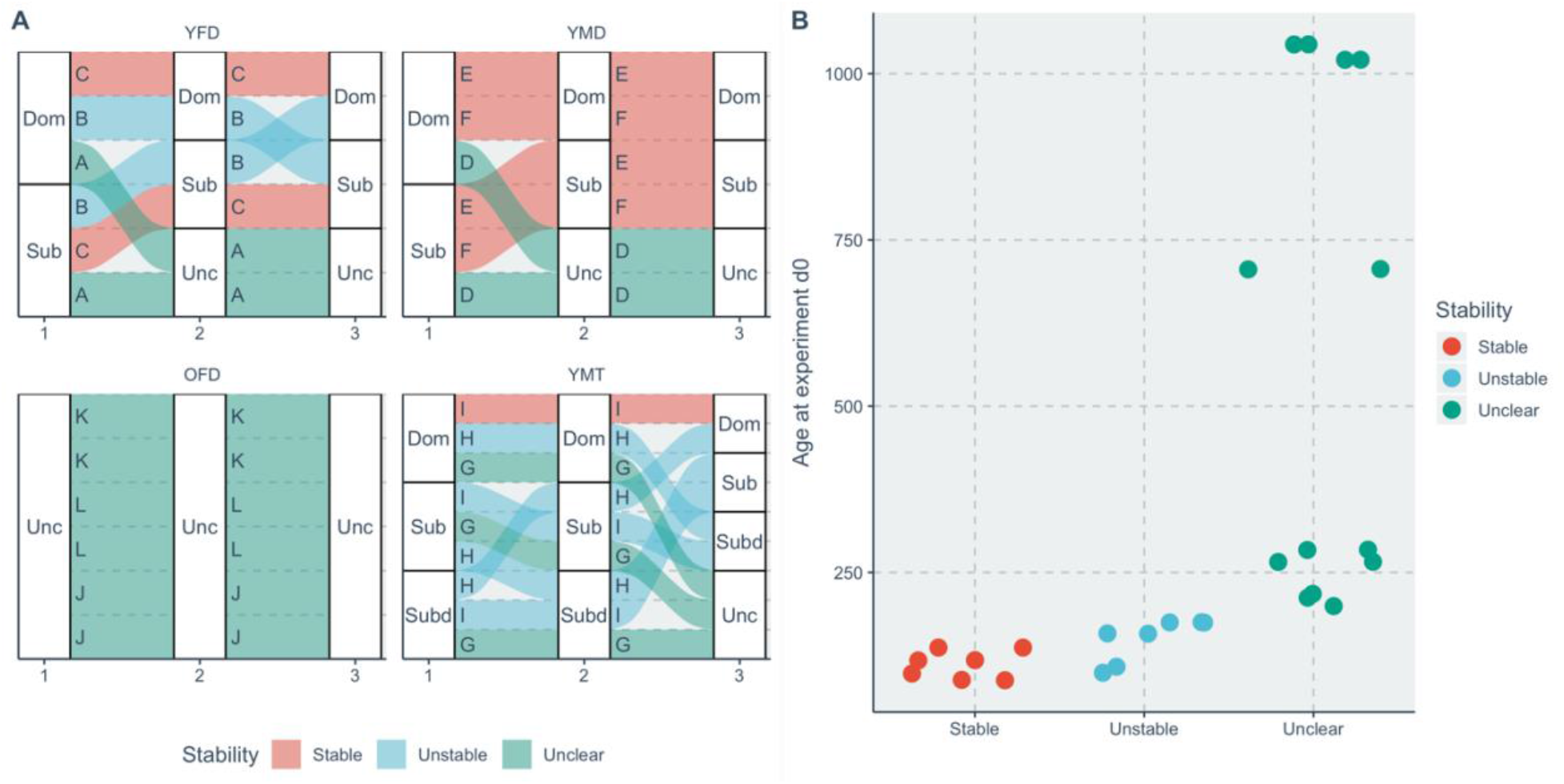
Social stability according to housing condition and across age. **A**: Alluvial or flow diagram showing the social dominance status switches across weeks, for each cage/individual labeled with a letter. *Note, no video data was recorded for the third week for cages J, K, or L due to the COVID-19 pandemic, but their stability was still considered “unclear” **B**: Scatter plot showing the relationship of age and social stability.

### Differences in the frequency of agonistic behavior types between sexes and male group-sizes, depending on dominance status

Next, differences were compared in the type of dominance behavior shown according to our ethogram (SI Table 1). A linear mixed model was used to compare within dominants, within subordinates, and across dominants and subordinates— subdominants were excluded from the analyses. For animals in dyads, animal-identity was used as a random factor while the behavior type and sex were used as fixed factors. When comparing the frequency of agonistic offensive behavior for dominant animals in dyads, the model determined that dominant animals significantly differed in the frequency of the type of behavior (Fig. 3A, Table 1), and a post-hoc comparison indicated that chasing was significantly more frequent than mounting, attacking, or food stealing (SI Table 2). There were no significant differences between the sexes, nor the interaction of behavior type and sex. When comparing subordinate agonistic defensive behavior, the model determined that subordinates in dyads also significantly differed in the frequency of the type of behavior (Fig. 3B, Table 3), and post-hoc comparisons indicated that they engaged in more fleeing than freezing (Estimate=74, SE=36.766, t=2.013, p=0.079), but there was no difference for induced flees (Estimate=36.333, SE=36.766, t=0.988, p=0.352). There were no significant differences between the sexes, nor the interaction of behavior type and sex. The total agonistic offensive behavior of dominants was also compared to the total defensive behavior of subordinates for dyads, using a mixed-model with cage-identity as a random factor. The model determined that there was no significant difference in the type of behavior, between the sexes, or the interaction between behavior type and sex (Table 3).

**Fig. 3:**
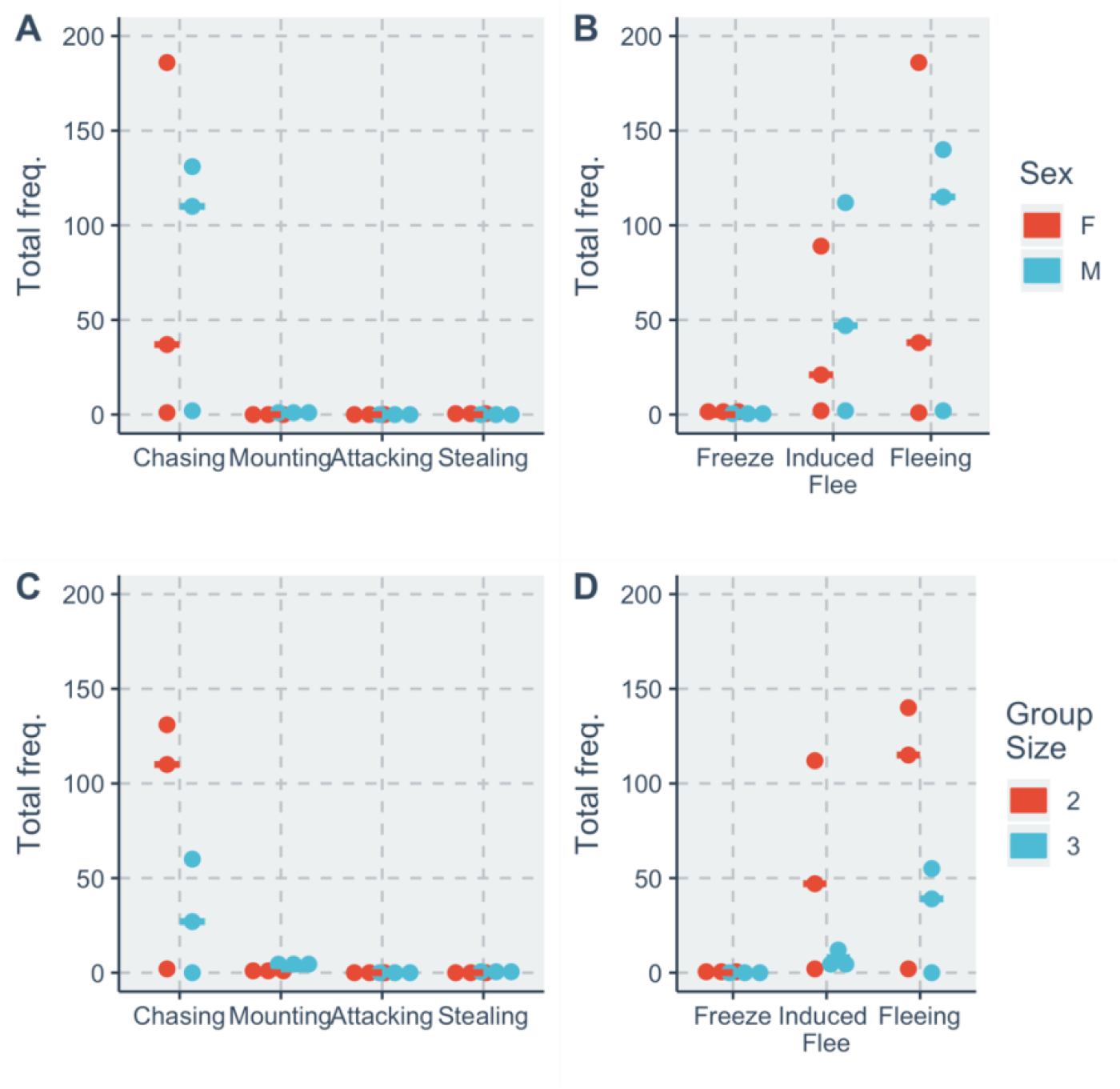
Frequency plots of agonistic behavior in dominant and subordinate *Acomys*. **A**: Scatterplot of agonistic offensive behaviors for dominant animals in dyads, with a horizontal bar denoting median **B**: Scatterplot of agonistic defensive behaviors for subordinate animals in dyads, with a horizontal bar denoting median condition. **C**: Scatterplot of agonistic offensive behaviors for dominant males in dyads or triads, with a horizontal bar denoting median condition. **D**: Scatterplot of agonistic defensive behaviors for subordinate males in dyads or triads, with a horizontal bar denoting median condition.

**Table 1:** Linear mixed effect models for estimating individual agonistic behavior

| Model | Fixed Effect | df | *F* | *p* |
| --- | --- | --- | --- | --- |
| Dominants in dyads,<br>agonistic offensive | **Behavior** | **3,12** | **5.042** | **0.017** |
|  | Sex | 1,4 | 0.009 | 0.928 |
|  | Behavior*Sex | 3,12 | 0.008 | 0.999 |
| Subordinates in dyads,<br>agonistic defensive | **Behavior** | **2,8** | **4.720** | **0.044** |
|  | Sex | 1,4 | 0.058 | 0.822 |
|  | Behavior*Sex | 2,8 | 0.055 | 0.946 |
| Dominants vs. Subordinates in<br>dyads | Off vs. Def | 1,4 | 5.052 | 0.088 |
|  | Sex | 1,4 | 0.034 | 0.863 |
|  | (Off vs. Def)*Sex | 1,4 | 0.211 | 0.670 |
| Male dominants,<br>agonistic offensive | **Behavior** | **3,12** | **6.203** | **0.009** |
|  | Group Size | 1,4 | 1.235 | 0.329 |
|  | Behavior*Group Size | 3,12 | 1.490 | 0.267 |
| Male subordinates, agonistic<br>defensive | **Behavior** | **2,8** | **5.512** | **0.031** |
|  | Group Size | 1,4 | 1.837 | 0.247 |
|  | Behavior*Group Size | 2,8 | 1.382 | 0.305 |
| Male Dominants vs. Subordinates | Off vs. Def | 1,4 | 3.369 | 0.140 |
|  | Group Size | 1,4 | 1.627 | 0.271 |
|  | (Off vs. Def)*Group Size | 1,4 | 2.330 | 0.202 |

A similar set of linear mixed models were used to determine differences between males in different group sizes. Again, the random effect was cage-identity, while the fixed effects were behavior type and group-size. Comparing dominant males in different group sizes, the model determined that they significantly differed in the type of agonistic offensive behavior (Fig. 3C, Table 3), and post-hoc comparisons indicated that chasing was the most frequent compared to all other behaviors (SI Table 3). There were no significant differences between the group sizes, nor the interaction of behavior type and group size. Comparing subordinate males in different group sizes, the model determined that they significantly differed in the type of agonistic defensive behavior (Fig. 3D, Table 1), and post-hoc comparisons indicated that fleeing was significantly more frequent than freezing (Estimate=85.333, SE=24.852, t=3.434, p=0.009), while there was no significant difference between freezing and induced fleeing (Estimate=53.333, SE=24.852, t=2.146, p=0.064). There were no significant differences between the group sizes, nor the interaction of behavior type and group size.

Comparing dominant and subordinate males, the model determined that there were no significant differences in their agonistic behaviors, between group sizes, or the interaction of behavior type and group size (Table 1).

### Differences in agonistic and huddling behavior during the dark cycle

Next, a separate dataset was used that collected the presence or absence of activity, chasing, induced flee, side huddle, and mounted huddle (see SI Table 4) per minute for the first 15-minutes for every hour in the dark cycle (20:00h to 06:00h). This data was collected for each cage, for three consecutive nights for three consecutive weeks (i.e., experimental days 1-3, 8-10, and 15-17). Here, differences in the proportion of time a behavior was present are compared between dyads of different sexes, males in different group sizes, and females of different ages. For each of the housing conditions, a generalized linear model was used with behavior, housing condition, and their interaction as fixed effects.

For male and female dyads, there was no significant difference between types of behaviors (*F*(3,20)=2.278, p=0.119), nor between sexes (*F*(1,19)=0.095, p=0.762), nor the interaction of sex and behavior type (*F*(3,16)=1.506, p=0.251). For males in different group sizes, there was a significant difference between types of behaviors (*F*(3,20)=7.759, p=0.002) (Fig. 4A), but not between the group sizes (*F*(1,19)=0.081, p=0.780), or the interaction of group size and behavior type (*F*(3,16)=0.187, p=0.904). Post-hoc comparisons indicated that Side huddle was the most frequent behavior (Estimate=0.156, SE=0.057, t=2.747, p=0.0143). For females in different age groups, there was a significant effect of behavior (*F*(3,20)=6.518, p=0.004) (Fig. 4B), with no significant difference between age groups (*F*(1,19)=0.002, p=0.969), but a significant interaction between age group and type of behavior (*F*(3,16)=3.621, p=0.036). Post-hoc comparisons indicated that the most frequent behaviors were mounted huddle (t=3.384, p=0.004) and side huddle (t=4.152, p=0.001). With young females engaging in significantly less side huddling than older females (t=-2.823, p=0.012).

**Fig. 4:**
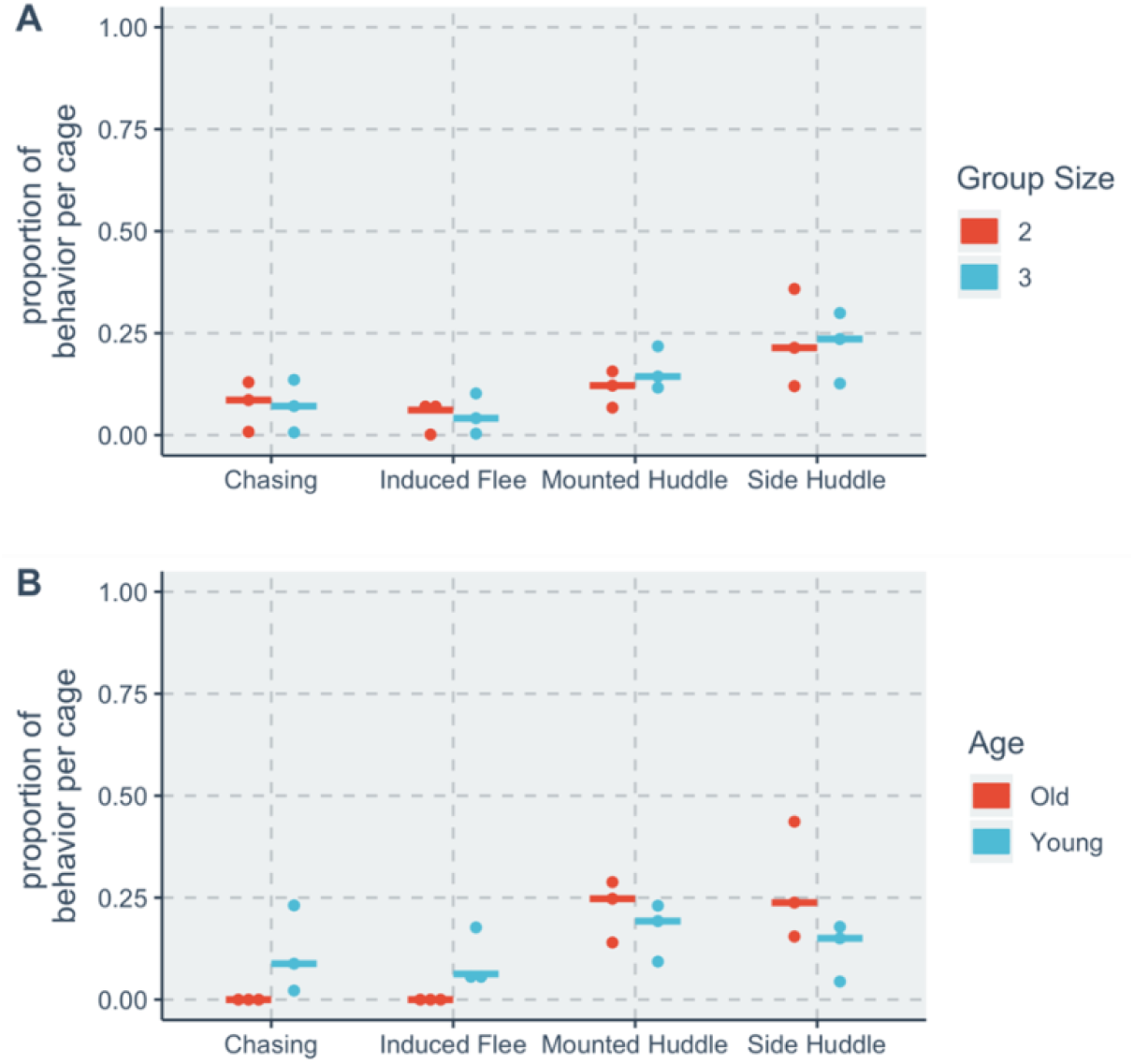
Proportion of behaviors throughout the night for cages of *Acomys*. **A**: Scatterplot comparing behaviors for males in different group sizes **B**: Scatterplot comparing behaviors for females with different ages.

### Differences in time to complete ear-hole regeneration

We then investigated whether differences in social dominance accounted for any phenotypic diversity in days to close the ear-hole after 4mm punch biopsy. A linear mixed model, with cage-identity as a random factor and David’s score and stability as fixed factors, predicted that David’s score had no significant effect on the days it took to close the ear-hole (*F*(1,9)=0.538, p=0.481), while there was a significant effect for social stability (*F*(2, 10)=9.997, p=0.008), and the interaction of David’s score and stability (*F*(2, 9)=8.620, p=0.032) (Fig. 5). Post-hoc comparisons indicated that those in unclear relationships took longer to regenerate than those in stable relationships (t=3.965, p=0.006), and that while dominants regenerated sooner in unclear groups, subordinates regenerated sooner in stable groups (t=-2.796, p=0.020). An exploratory analysis of the data, however, indicated that age played a significant role in the time to complete ear-hole regeneration (*F*(1,8)=4.028, p=0.082), and removed the effect of stability when entered into the linear mixed effect model (*F*(2,10)=0.022, p=0.979), but not the interaction of David’s score and stability (*F*(2,9)=5.362, p=0.031) (SI Fig. 1).

**Fig. 5:**
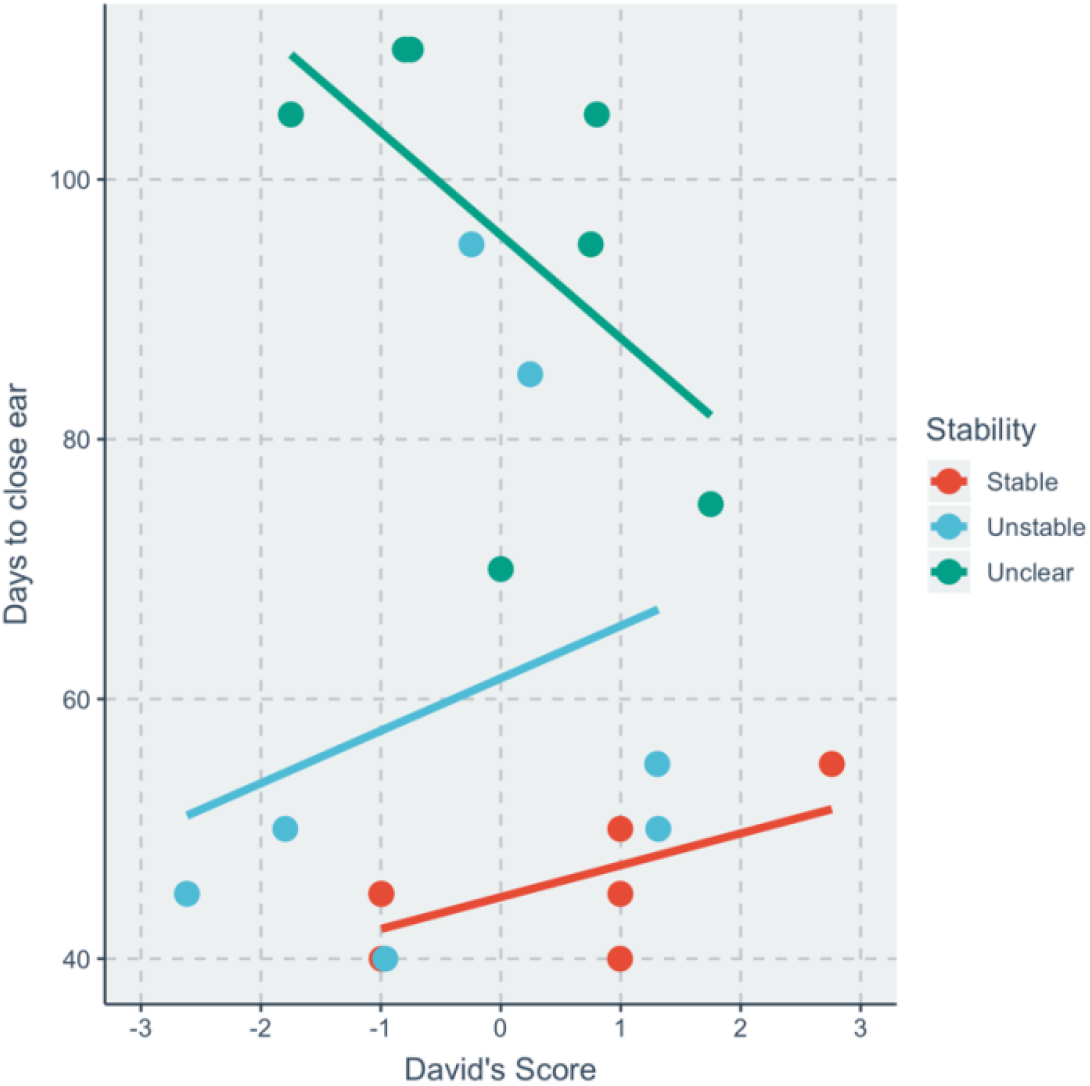
Time to close the ear-hole after injury in relation to David’s Score and social stability. Scatterplot of all animals with estimated line reflecting the linear mixed effects model.

### Blood cortisol

Differences in blood serum cortisol were compared in relation to David’s score and stability, as well as whether blood cortisol correlated with time to close the ear-hole. Only males were included in the analysis since females consistently had high values that went beyond the detectable limit of the assay. A linear mixed effect model, with cage-identity as a random factor and David’s score and stability as fixed factors, predicted that David’s score had no significant effect on the blood cortisol concentration of males (*F*(1,7)=0.022, p=0.887), with stability status also having no significant effect (*F*(2, 6)=0.516, p=0.623), nor the interaction of David’s score and stability (*F*(2,6)=3.070, p=0.114). A simple Spearman correlation determined there was no significant correlation between the time to close the ear-hole and blood cortisol levels (r=-0.299, p=0.279).

### Cortisol synthesis genes

Differences in genes expression ratios required for cortisol synthesis in the adrenal gland were then compared depending on sex, David’s score, and social stability using RTqPCR. Linear mixed effects models with cage as a random effect and sex, David’s score, and social stability as fixed effects indicated that only Nr5a1 significantly differed for the interaction of David’s score and stability (*F*(2,13)=4.014, p=0.043) (Fig. 6), while all other factors had no significant effect on gene expression ratio for each gene tested (i.e., Cyp11a1, Nr5a1, Nrb01, and StAR) (SI Table 5). Post-hoc comparisons for Nr5a1, a transcriptional regulator of StAR, indicated that as David’s score increased for unclear groups, Nr5a1 significantly decreased compared to a significant increase for stable groups (t=-2.793, p=0.015).

**Fig. 6:**
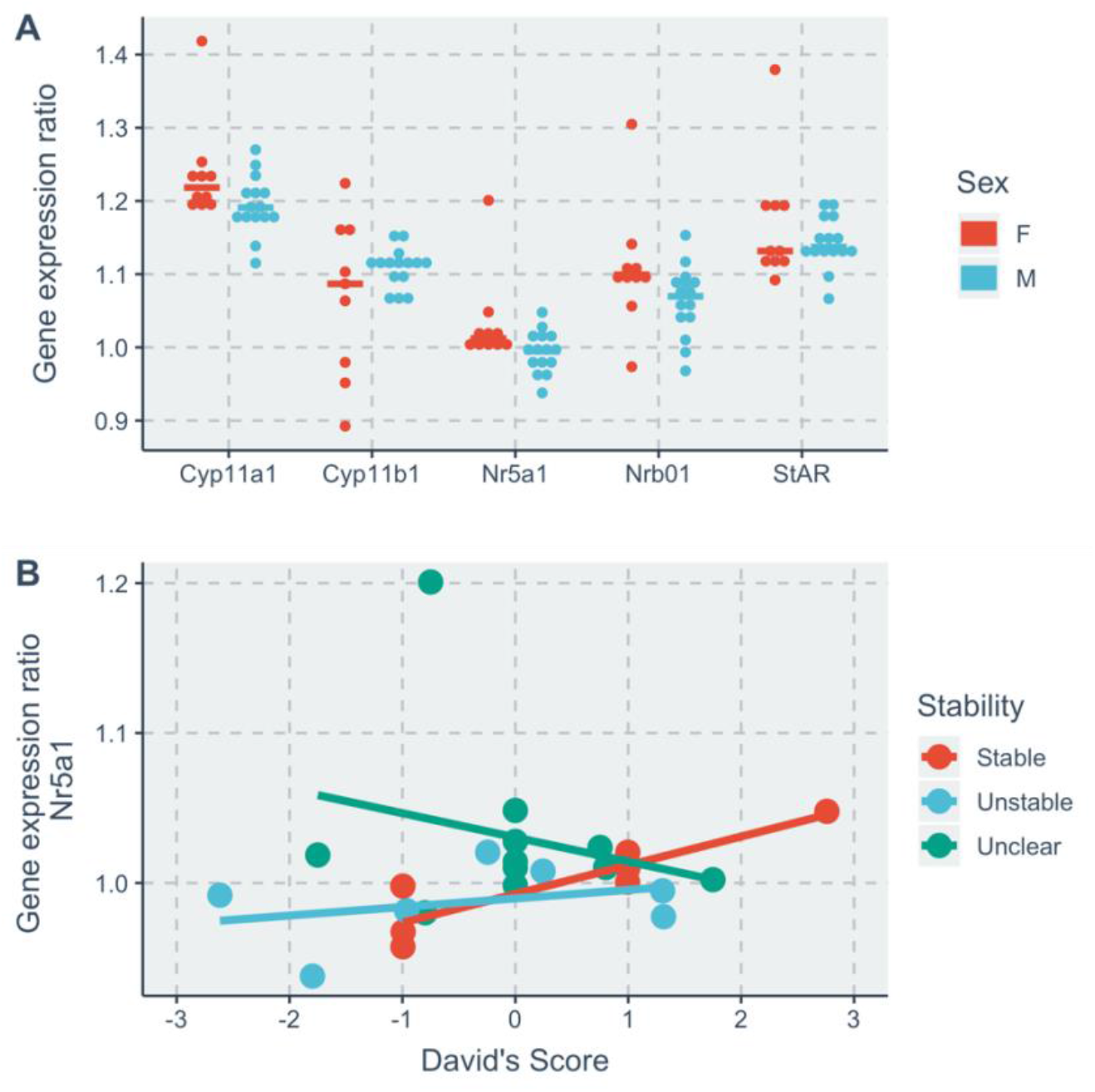
Gene expression ratios according to sex or David’s score and social stability. **A**: Scatterplot of gene expression ratio for all cortisol synthesis genes tested comparing sexes, individual points for each animal. **B**: Scatterplot of gene expression ratios for Nr5a1 in relation to David’s Score and social stability, points denote animals while lines denote linear mixed effects model.

## Discussion

The primary aim of this study was to investigate the stability of social dominance relationships in groups of adult *Acomys*, investigate any sex differences, and determine what behaviors they may engage in rather than freezing. A secondary aim was to investigate phenotypic diversity related to social dominance and stability in measures of ear-hole closure, blood cortisol, and genes related to cortisol synthesis. Overall, the results indicate that most individuals in a group could be assigned a social dominance status, but most groups rarely engaged in agonistic behavior making the assessment of social stability unclear. There were no significant effects between the sexes in agonistic behavior. As predicted, *Acomys* rarely froze in response to an agonistic offensive behavior, and more often fled or induced fled (e.g., were displaced). They also more frequently huddled during their active cycle than engaged in agonistic behavior. Secondary investigations into phenotypic diversity determined that social stability was associated with slower ear-hole closure, but this was modified by differences in age. Females had higher levels of blood cortisol but were excluded from analyses because their levels went beyond the limits of the assay. Regarding cortisol synthesis, there was a significant effect for the interaction of David’s score and social stability for a transcriptional enhancer of StAR, Nr5a1. This result however should be taken with caution, as further discussed.

### Agonistic behavior and social dominance relationships

The current study reproduced the effects reported in the initial study on agonistic behavior in *Acomys* (38), and expanded our understanding of social dominance status and social stability. The main findings by Porter (38) were that males and females both engage in overt and offensive agonistic behavior like chasing and attacking, that they rarely freeze or show “appeasement gesture(s)”, and that females are often dominant over males. As expected, the current study also found no significant differences between the sexes in agonistic offensive or defensive behavior, and that *Acomys* rarely froze in response to agonistic offensive behavior—more often fleeing or induced fleeing (defined by Porter as displacing but see further discussion). While the current study excluded mixed-sex groups, it systematically recorded individual agonistic interactions across three weeks to determine dominance rankings and their stability—an aspect missing from Porter’s initial studies. These repeated recordings of agonistic behavior determined that most groups had clear dominance rankings of dominant, subdominant, or subordinate (where appropriate given group size). However, many groups did not engage in agonistic behavior throughout the observation time, making the stability of their social dominance relationships unclear. Of the groups that consistently engaged in agonistic behavior, just as many were stable as unstable. This adds to the limited literature on agonistic behavior in *Acomys* by reproducing main findings from a classic study and expanding information on their social dominance relationships.

While a lack of freezing or subordinate posturing in mice and rats often coincides with unstable dominance relationships (2,3), the lack of freezing in *Acomys* coincided with low levels of agonistic behavior and stable relationships. This finding contradicts our general understanding of social dominance relationships in Murids (e.g., mice and rats) and indicates that *Acomys* are likely engaging in other behaviors beyond freezing or subordinate posturing. Currently the alternative behaviors to freezing or subordinate posturing remain unclear.

Some studies on wild-mice in larger housing (∼18,580cm^2^) suggest that they often avoid one another, termed as spatial segregation (54). Thus, a study comparing larger housing to standard housing while comparing subordinate and segregation behaviors between *Acomys* and common rodents (e.g., mice and rats) would help elucidate whether *Acomys* favor avoidance behaviors over freezing. We attempted to study spatial segregation in the current study (*unreported*) but were limited by the relatively small cage size (1800cm^2^)—albeit it was much larger than standard mouse cages (∼430cm^2^ to 500cm^2^). This relatively small cage size made it difficult to clearly delineate points of spatial segregation, leading us to define displacements (as defined by Porter (38)) as induced flees because it was unclear whether the dominant animal was occupying the space previously occupied by the subordinate.

Another important follow-up study should consider the formation of dominance relationships. A recent study on the temporal microstructure of dyadic social behaviors during relationship formation in mice (44) revealed that tail-rattling may deescalate aggressive behavior in pairs, and dominants increase allogrooming while subordinates were more likely to avoid the head of the dominant while investigating or huddling. This provides great insight into how a dominance relationship is formed in mice and similar studies in *Acomys* may provide key details on latent “appeasement gesture(s)”, and species differences in agonistic behaviors.

Given our lack of agonistic behavior in some groups, particularly old females, it would also be fruitful to devise social dominance paradigms beyond home-cage behavior.

Briefly, other studies measuring social dominance in rodents use a diverse set of methods to measure dyadic social relationships (22) through social conflict in a narrow tube (55,56) or competing for rewards (57), for example. While these tests are not guaranteed to correlate with home-cage behavior or one another (58,59), they provide further insight into the social relationship and can be used in a round-robin tournament to measure dominance relationships with equal numbers of social interactions.

### Phenotypic diversity in ear-hole closure

In addition to determining the social dominance status of individual animals and the stability of their status within the group, the current study also measured phenotypic differences in ear-hole closure after injury and glucocorticoids. As expected, all the animals closed their ear-holes, indicating that their ear tissue regenerated after injury (28–30). However, there was great diversity in the time to close the ear-hole ranging from 40 to 110 days. Some of this diversity could be accounted for by (i) social stability, with *Acomys* in unclear relationships taking longer to regenerate, and by (ii) the interaction of an individual’s David’s score and the social stability, where individuals with higher David’s scores regenerated quicker in unclear groups while those with higher David’s scores in stable groups regenerated slower. However, differences in stability were highly correlated with differences in age, and when age was added to the statistical model, the significant effect of stability was reduced to a non-significant effect. This was because the older animals were more likely to have an unclear dominance relationship. Thus, the current study cannot adjudicate on whether variability in ear-hole closure is likely due to the social dominance context or age.

Other studies find that age coincides with slower or faster regeneration (60–62) but also find other secondary factors (e.g., nutrition, seasonal variation, or stress) are associated with slower regeneration (63,64). A small study in *Acomys* found that older *Acomys* (≥3 years) regenerated 2mm biopsy punches to the ears slower than younger *Acomys* (2-months) (65). However, this was only a 1-week difference, which starkly contrasts with the 10-week range of the current study. Notably, this wide range isn’t unique to our study. Other studies measuring ear-hole closure in *Acomys* have also found great diversity in the time to close the ear-hole (ranging from 20 to 90 days, Supplement table 6) (28–31). Some post-hoc exploratory analyses in those studies suggest that other factors like blood draws and lactation likely contributed to this variability (30). The current study, however, restricted blood collection to post-mortem and no breeding animals were used. Thus, other factors beyond aging, blood draws, and lactation are likely contributing to variability in *Acomys* ear-hole closure.

Overall, it is likely that older *Acomys* may significantly differ in their rate of tissue regeneration, and that the effects we see with dominance are just correlated with age. Indeed, animals that have been housed together longer should have lower levels of agonistic behavior (2,3). Moreover, regenerative ability is known to decline with age in many animals (60,61). Future studies on *Acomys* regeneration should thus consider the role of age, and a systematic study and other latent factors relating to heredity, development, and the environment.

### Phenotypic diversity in glucocorticoids

The hypothesis regarding social dominance and regeneration was that subordinate animals and those in unstable groups would have higher basal levels of cortisol, and slower regeneration due to the combination of increased stress hormones and energetic demands of their social environment (10,34,35,66). Unfortunately, our data do not support this hypothesis. However, it is important to note, these serum hormone and gene expression data were derived from resting, non-stressed animals after the wounding and healing were complete. Thus, future studies would be needed to fully address the role that HPA activity and reactivity may play in mediating the relationship between social stability, dominance status, and regeneration. Moreover, social relationships can also be positive and reduce glucocorticoid levels, thereby making it difficult to determine how social dominance relationships affect HPA activity and reactivity (7,67–69).

Despite these limitations we did however find that blood cortisol levels were lower for males than females (albeit it was not possible to statistically compare them), but there were no significant differences between the sexes in the expression of genes for cortisol synthesis that were measured. Others have also found a sex difference in resting cortisol levels in *Acomys*, with females having substantially more cortisol than males (50,70,71). Thus, it is likely that the observed sex difference is representative of *Acomys* in general and other factors may be responsible for this difference beyond the cortisol synthesis genes tested.

There were no significant differences in basal blood serum cortisol levels between *Acomys* cage-mates with different David’s scores or between stable, unstable, or unclear cages. Moreover, differences in cortisol levels did not significantly correlate with variability in time to close the ear hole after injury. Some limited evidence comparing genes involved in adrenal cortisol synthesis indicated that the gene Nr5a1 was differentially expressed regarding the interaction between David’s scores and stability. Nr5a1 is a transcriptional enhancer of StAR, which makes pregnenolone from cholesterol—the first step in making cortisol from cholesterol. Without further differences in blood cortisol levels, however, it is unclear what role it may have regarding social dominance or ear-hole regeneration.

### Conclusion

Overall we found that both female and male *Acomys* readily engage in agonistic behavior, but more often huddle during the active dark cycle—especially older females. Most individuals in a group have a social dominance status, but some groups are unstable or have unclear statuses because of their infrequent agonistic behavior. As observed in previous studies, *Acomys* rarely froze in agonistic interactions and more often fled, significantly. Previous studies report subordinate animals heal wounds slower (11), but we found little evidence for a similar phenomenon in *Acomys* tissue regeneration. Rather, there was slower wound healing in older animals. Studies also suggest that subordinate animals and those in unstable groups have increased glucocorticoids, but we found little evidence for this in *Acomys*. Future studies should consider the limitations discussed for this study and also consider more empirical investigation of social dominance by experimentally manipulating social relationships via paradigms like social defeat stress (72), resident intruder paradigms (73), chronic subordinate housing (74), or controlled physical injury (75), which can all modify immune responses or wound healing. This might provide more insight into how the social relationships of *Acomys* may mask treatment effects measuring wound healing and glucocorticoids.

## Methods

### Experimental design

There was great heterogeneity in group-size (i.e., dyads and trios), age (i.e., 12-41 weeks old and 100-149 weeks old), and sex of the animals available for this study. This allowed us to make several comparisons: The effect of sex comparing adult (12-41 weeks old) male and female dyads (YMD vs YFD, n of 3 per experimental group), the effect of group-size in males comparing adult (12-41 weeks old) male dyads and trios (YMD vs YMT, n of 3 per experimental group), and the effect of age in female dyads comparing young (12-31 weeks old) and old female (100-149 weeks old) dyads (YFD vs OFD, n of 3 per experimental group). In a more balanced study (i.e., a factorial design), we would also have trios of females and/or old-age male dyads, but they were rare in our colony for the duration of this study.

A total of 15 male and 12 female *Acomys cahirinus* bred at the University of Florida were housed together since weaning at 21 to 41 days and in adulthood the following measures were collected: (a) general incidence of agonistic and huddling behavior during the dark cycle, (b) dominance and avoidance behavior during two 10-minute recordings at the start of the night, (c) time to regenerate ear-tissue after biopsy punch, (d) blood serum cortisol levels with radioimmunoassay, and (e) quantitative mRNA levels of cortisol synthesis genes from the adrenal gland with RT-qPCR (see SI Fig. 2 for an experimental timeline).

### Attrition

Due to the COVID-19 pandemic, there was some data attrition for this experiment. Specifically, for the old females in dyads (OFD) there was no home-cage behavior, or ear-hole regeneration data collected during the third week of recordings for cages J, K, and L. We did however collect blood and adrenal tissue for the glucocorticoid measures for cages J and K, but not for cage L.

### General husbandry procedures

All *Acomys* were kept on a 14:10 light/dark cycle with lights on at 06:00. A red-light emitting diode (LED) was on at the start of the dark cycle 20:00-24:00 to observe the animals during dominance rank observation, while infra-red lighting remained on throughout the entire dark cycle. The temperature was 27 ± 3ºC and humidity around 50%. Animals were housed in either Techniplast GR1800 double-decker cages (floor area: 1862cm^2^) (10 females, and 5 males) or NexGen Rat 1800 cages with two levels (floor area: 1800cm^2^) (2 females, 10 males). This difference in housing was because the Techniplast cages started to break and were not ideal for handling *Acomys*, thus the NexGen cages replaced the Techniplast cages in our colony. All cages contained aspen wood chip bedding, 2cm deep, and animals had *ad libitum* access to standard chow (Teklad 2918) and tap water, with food supplementation one day per week as determined by a veterinarian (e.g., grapes, mealworms, carrots, broccoli, sweet potatoes). *Acomys* were provided with two shelters (Bio-Serv, red Rat Retreats), a nylabone, a wood gnawing block, and a chewing stick. All animals were individually identified by a small hair-shave to the fore-limb or hind-limb.

### Collection of social dominance behavior

Behavioral repertories of agonistic behavior were video-recorded in the home-cage for two 10-minute periods each day for three subsequent days(76) for three weeks (experiment days 1-3, 8-10, and 15-17) during the first three hours of the dark cycle (20:00h to 23:00h)—exclusive collection during the dark cycle was determined by 72-hour screening of activity (See SI Text 1 and SI Fig. 3). Individual agonistic behavior was coded (i.e., focal sampling) in BORIS for the frequency and duration of all occurrences during the 10-minute period using our agonistic behavior ethogram (SI Table 1). Individuals were identified by a combination of fur-shavings to the upper portion of their limbs and/or tail length (since *Acomys* are prone to losing their tails(77)). Inter-rater reliability was good (10% sub-sample, mean kappa 0.87), as was intra-rater reliability (15% sub-sample, mean kappa 0.96).

### Calculation of social dominance status, directional consistency index, and stability

From the collection of social dominance behavior we calculated individual social dominance status, a group-level directional consistency index, and individual dominance status stability categorizations. Social dominance status was calculated using a David’s score, which is an index of the proportion of wins adjusted for the strengths of their opponents (i.e., how often their opponents win or lose) (78). David’s scores were then ranked from highest to lowest within cage-groups, and the animal with the highest score was determined as dominant, the lowest score as subordinate, and the middle score (if applicable) as subdominant. Animals that did not engage in agonistic behavior and had a David’s score of 0 were categorized as having an unclear social dominance status. A directional consistency (dc) index was then calculated which measures the degree of the directionality of agonistic behaviors from the most dominant towards the most subordinate(79). An index score of 1 indicates that all offensive agonistic interactions were in the direction of the most dominant to the most subordinate, while a score of 0 indicates that there are equal numbers of offensive agonistic interactions between the pair. Index scores were averaged for groups of three to be more comparable to the scores for groups of two. Finally, stability of social dominance was determined by calculating the social dominance status from the David’s score for each week and then categorizing individuals as stable or unstable (4,21).

Those that maintained the same social dominance status across all weeks were categorized as stable, and those that switched social dominance status were designated as unstable. All dominance calculations were completed using the ‘compete’ package v0.1(80) in R (v4.2.0).

### Collection of group agonistic and huddling behavior

Using the same video-recording and one/zero sampling methods described in SI Text 1, we measured differences in group agonistic and huddling behavior using our specific ethogram for all animals (SI Table 4). Inter-rater reliability was good (10% sub-sample, Mean kappa of 0.96), as was intra-rater reliability (5% sub-sample, Mean kappa of 0.92). Video observations with a kappa value of 0.85 were coded a second time and discussed. If the cage was not clearly visible during coding, another 15-minute period during the respective hour was chosen (e.g., minutes 30 to 45 or 44 to 59—this only occurred during the hours of 21:00h and 23:00h when the social dominance recordings were collected).

### Ear-punch biopsy injury and regeneration

Around experimental day 22, Animals were anesthetized with 4% (v/v) vaporized isoflurane (Pivetal®, Patterson Vet Supply) and administered a through-and-through hole in either the left or right ear pinna using a 4-mm biopsy punch (Robbins Instruments, Chatham, NJ) ∼1mm distal from the head and centered on the middle of the pinna. Following injury, the animals were lightly anesthetized for several minutes every 5 days until the hole was closed for measurements. Measurements were made using calipers by measuring the diameter of the proximal-distal (DPD) and the anterior-posterior (DAP) axes for each ear hole. The ear-hole area was calculated for an ellipse to account for unevenness across the hole (equation 1). When no light could be seen through the hole in the ear, it was considered closed. Experimenters were blinded to dominance status calculations.

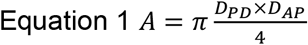

### Blood collection and cortisol analyses

Animals were euthanized several days after their ear-hole injuries were closed. Immediately following euthanasia by carbon dioxide asphyxiation, animals were decapitated, and trunk blood was collected in clean 1.2ml eppendorf tubes and placed on the bench at room temperature for ∼3minutes to allow the blood to clot. The tubes were then spun down in the centrifuge and the serum supernatant was collected and stored in the freezer at −20ºC. Serum cortisol concentrations were measured using commercially available radioimmunoassay kits (MP Biomedicals; Solon, OH) and performed as indicated by the supplier (catalog # 07-221102R). Samples were run in duplicate and values averaged. All duplicate samples had a coefficient of variation (CV) under 10%. The intra-assay CV was 11.05% and the lowest and highest detectable values were 9.5 and 1019 ng/mL, respectively. Experimenters were blinded to sex, and dominance calculations.

### Adrenal organ collection and mRNA expression

Immediately following trunk blood collection, both the left and right adrenal glands were harvested from the animals and immersed in RNAlater (Invitrogen) at 4ºC for 24-hours and then stored in the freezer at −80ºC. RNA was isolated from both adrenal glands together using the RNAeasy mini kit (Qiagen) following the manufactures recommended protocol, with tissue homogenization being performed using a rotor stator type tissue homogenizer (ProScientific Bio-Gen PRO200 Homogenizer; Multi-Gen 7XL Generator Probes) in RLT Buffer (Qiagen). RNA quality was assayed using a nanodrop. The cDNA was generated from 500ng of RNA using ezDNase Enzyme (Invitrogen) and SuperScript IV VILO Master Mix (Invitrogen) following the manufacture’s protocol (81). RT-qPCR was performed using Sso-Fast EvaGreen Supermix (Bio-Rad) on a Bio-Rad C1000 Touch Thermal Cycler in triplicate for each sample (n=21). The fold change in gene expression was calculated accounting for primer efficiency and using the Pfaffl method (82) with ActinB as the reference gene. The sequence of Acomys-specific PCR primers designed from a preliminary genome can be found in SI Table 7. All reactions were run with an annealing temperature of 60ºC. Experimenters were blinded to sex, dominance rank, and stability.

### Statistical analyses

All statistical analyses were performed in R (v4.2.0) and used p<0.05 as the critical threshold. The normality and homogeneity of variance in each dataset were examined graphically, and no transformations were performed. Linear mixed-models were run as described in the results with the lmerTest package (v3.1.3), while pair-wise comparisons were calculated by hand following the methods of Siegel and Castellan (83).

## Ethics

This study was carried out in accordance with the University of Florida IACUC protocol (201807707).

## Data, code, and materials

All code and additional materials can be found at the following github repository: https://github.com/javarhol/AcomysDominance_2022_Data_Results.git The data will be provided by the corresponding author upon email request.

## Competing interests

The authors declare no competing interests

## Acknowledgments

The authors gratefully acknowledge the animal care staff and veterinarians at the University of Florida. The research was supported by the SNSF Early Postdoc Mobility Fellowship P2BEP3_181707 awarded to J.A.V., a NIH T32 DK074367-14 supporting J.A.V., and a W.M. Keck Foundation grant awarded to M.M.

## Contributions

This research was conceptualized, supervised, and funding was acquired by J.A.V. and M.M.; Methodology by J.A.V, M.M, R.D.R., J.C., W.B.B.; Investigation by J.A.V., G.G., A.J., S.M., R.D.R., and M.M.; Resources by M.M., J.A.V., R.D.R. and W.B.B.; Software, Validation, Formal analysis, data curation, original draft, visualization, and project administration by J.A.V, and review and editing by all authors.

## Notes

https://github.com/javarhol/AcomysDominance_2022_Data_Results.git

### Competing Interest Statement

The authors have declared no competing interest.

