## Supplemental Information for "Social dominance status and social stability in spiny mice (*Acomys cahirinus*) and its relation to ear-hole regeneration and glucocorticoids"

[1]Department of Biology, University of Florida, Gainesville, FL, USA; [2]Department of Molecular Genetics and Microbiology, University of Florida, Gainesville, FL, USA; [3]Department of Psychology, University of Florida, Gainesville, FL, USA; [4]Department of Psychology and Neuroscience and Behavior, Barnard College of Columbia University, New York, NY, USA; [5]Genetics Institute, University of Florida, Gainesville, FL, USA

SI Table 1: Individual level social dominance behavior ethogram

| Behavior category | Behavior | Description |
|---|---|---|
| Offensive Agonistic Behavior | Chasing | Actor mouse rapidly pursues target mouse wile target mouse flees, unaccompanied by mounting |
| | Mounting | Actor mouse rapidly pursues target mouse wile target mouse flees, accompanied by side or rear mounting |
| | Attacking | Actor mouse lunges and/or bites target mouse, without target mouse counter-attacking |
| | Food stealing | Target mouse has control of food by holding it with both forepaws and/or mouth. Actor mouse then takes control over food, leaving target mouse without control |
| Defensive Agonistic Behavior | Flee | The actor mouse approaches the target mouse and the target mouse rapidly flees; the actor follows via chasing or mounting |
| | Induced Flee | The actor mouse approaches the target mouse, and the target rapidly flees; the actor does not follow |
| | Freeze | Target mouse is immobile in response to agonistic offensive behavior of actor mouse |
| Other | Inactive | Mouse visible and motionless for >15-seconds |
| | Unseen | Cannot reliably code whether mouse is in or on a shelter |
| Location | Top Left | Focal mouse is on the upper level on the left half of the cage |
| | Top Right | Focal mouse is on the upper level on the right half of the cage |
| | Bottom Left | Focal mouse is on the bedding level on the left half of the cage |
| | Bottom Right | Focal mouse is on the bedding level on the right half of the cage |

SI Table 2: Linear mixed effect model t-tests dyads, group agonistic behavior

| Model | Fixed effects | Estimate | Std. Error | t | p |
|---|---|---|---|---|---|
| Dominant ag bhv | Intercept | 74.667 | 24.506 | 3.047 | 0.008 |
| | Mounting | -74.667 | 34.589 | -2.159 | 0.052 |
| | Attacking | -74.667 | 34.589 | -2.159 | 0.052 |
| | Stealing | -74.333 | 34.589 | -2.149 | 0.053 |

SI Table 3: Linear mixed effect model t-tests males, group agonistic behavior

| Model | Fixed effects | Estimate | Std. Error | t | p |
|---|---|---|---|---|---|
| Dominant ag bhv | Intercept | 81.00 | 15.42 | 5.251 | 0.001 |
| | Mounting | -80.33 | 21.79 | -3.687 | 0.003 |
| | Attacking | -81.00 | 21.79 | -3.718 | 0.003 |
| | Stealing | -81.00 | 21.79 | -3.718 | 0.003 |

SI Table 4: Group-level agonistic and huddling behavior ethogram

| Behavior | Description |
|---|---|
| Activity | Animal is observable and performing any activity for at least 5-seconds; must see entire head of mouse |
| Chasing | The actor mouse approaches the target mouse and then rapidly follows the target mouse while it rapidly flees |
| Induced Flee | The actor mouse approaches the target mouse and the target mouse rapidly flees; the actor does not follow, it keeps a constant pace of activity throughout |
| Mounting (active) | The actor mouse approaches the target mouse and places its forepaws on the dorsal side of the target mouse; leading to a chase |
| Side-Huddle | One mouse is resting in direct body contact side-by-side |
| Mounted-Huddle | One mouse is resting with at least their forepaws on the dorsal side of the other mouse |

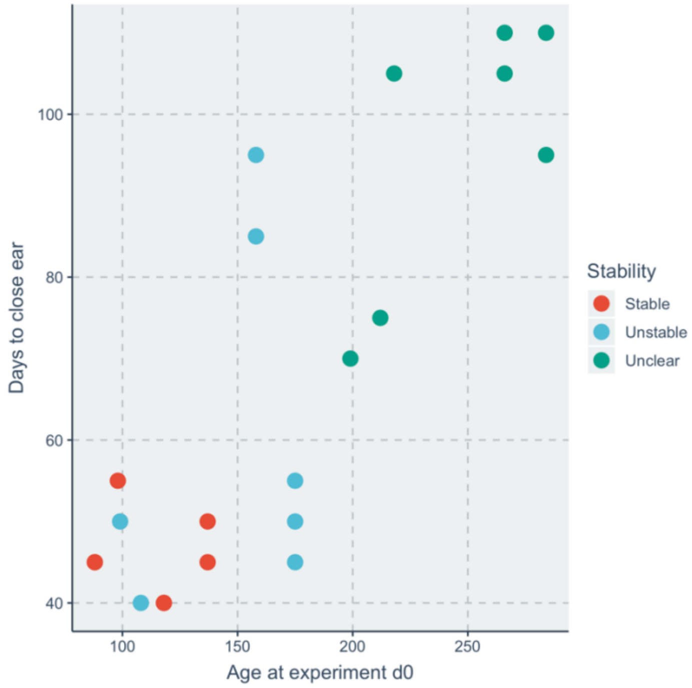


*SI Fig 1*: <u>The relationship of Age and Days to Close ear-hole.</u> Scatterplot showing the relationship between Age and Days to Close ear, with individuals colored by social stability.

SI Table 5: Linear mixed effect models for gene expression ratios

| Model | Fixed Effect | df | F | p |
|---|---|---|---|---|
| Cyp11a1 | Sex | 1,7 | 4.133 | 0.081 |
| | David's Score | 1,12 | 0.021 | 0.885 |
| | Stability | 2,7 | 0.036 | 0.965 |
| | David's Score*Stability | 2,14 | 2.081 | 0.161 |
| Cyp11b1 | Sex | 1,7 | 0.522 | 0.493 |
| | David's Score | 1,11 | 0.065 | 0.804 |
| | Stability | 2,11 | 0.686 | 0.524 |
| | David's Score*Stability | 2,11 | 0.177 | 0.840 |
| Nr5a1 | Sex | 1,7 | 1.917 | 0.208 |
| | David's Score | 1,12 | 0.036 | 0.852 |
| | Stability | 2,8 | 0.250 | 0.784 |
| | **David's Score*Stability** | **2,13** | **4.014** | **0.044** |
| Nrb01 | Sex | 1,6 | 3.422 | 0.115 |
| | David's Score | 1,11 | 0.169 | 0.689 |
| | Stability | 2,6 | 1.515 | 0.291 |
| | David's Score*Stability | 2,15 | 2.091 | 0.159 |
| StAR | Sex | 1,7 | 0.513 | 0.497 |
| | David's Score | 1,12 | 0.088 | 0.772 |
| | Stability | 2,8 | 0.132 | 0.879 |
| | David's Score*Stability | 2,13 | 1.019 | 0.388 |

SI Table 6: Ranges of ear-hole closure time for *Acomys* across published studies

| **Study** | **Species** | **Time to ear-hole closure** | **N** |
|---|---|---|---|
| Seifert et al., 2012 | *Acomys kempi* (wild) | 20 to 50 days | NA |
| Gawriluk et al., 2016 | *Acomys cahirinus* (lab) | 30 to 90 days | 40 |
| | *Acomys kempi* (wild) | 40 to 90 days | 15 |
| Matias Santos et al., 2016 | *Acomys cahirinus* (lab) | 25 to 55 days | 15 |
| Simkin et al., 2017 | *Acomys cahirinus* (lab) | 34 days | 6 |
| Current study | *Acomys cahirinus* (lab) | 40 to 110 days | 21 |

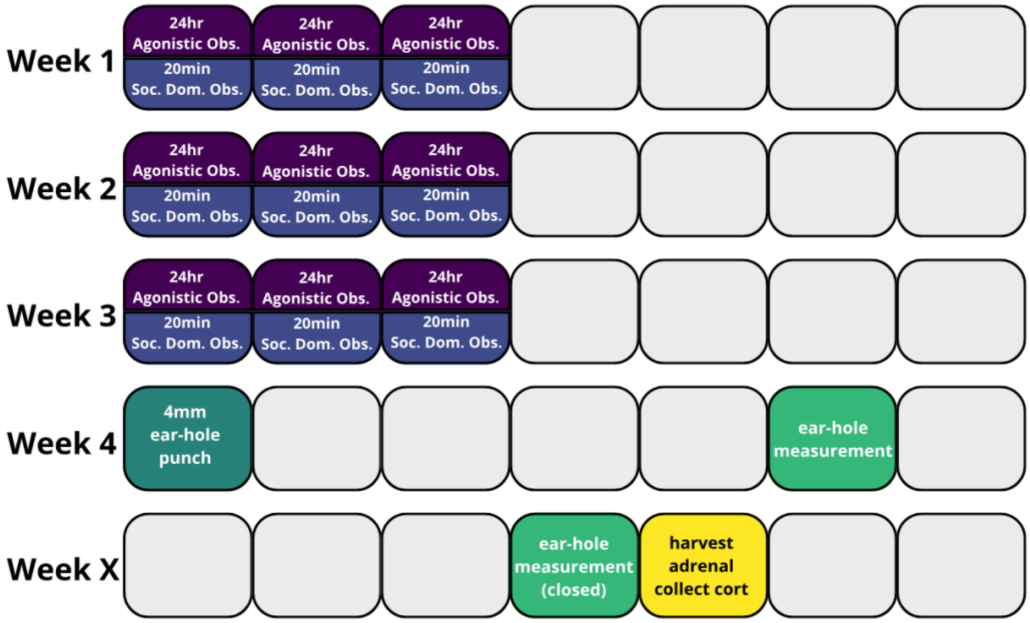


*SI Fig. 2:* Experimental timeline per week. Week X indicates that the time of harvesting adrenal gland and blood collection was dependent on the time it took to finish closing the ear-hole.

*SI Text 1: Determination of when to collect behavioral data*

To determine when the animals were engaging in agonistic behavior, we screened two groups of females and two groups of males for general activity and agonistic behavior (SI Table 4). Activity, or the animal moving for more than 5-seconds, was video recorded in the home-cage for three 24-hour periods once a week for three consecutive weeks (experiment days 1-3, 8-10, and 15-17). Recordings always began on the day of cage-change. Behaviors were coded in Behavioral Observation Research Interactive Software, BORIS (v.7.0+) using one/zero sampling of each 1-minute interval for the first 15-minutes of every hour [49] of the 24-hours for the 9 days of observation. Video coders were blinded to sex and social dominance rank of the animals. Here we found that the animals were most active during the dark cycle (20:00h-06:00h) and typically engaged in agonistic behavior around the start of the dark cycle at 21:00h (see supplementary Fig. 3).

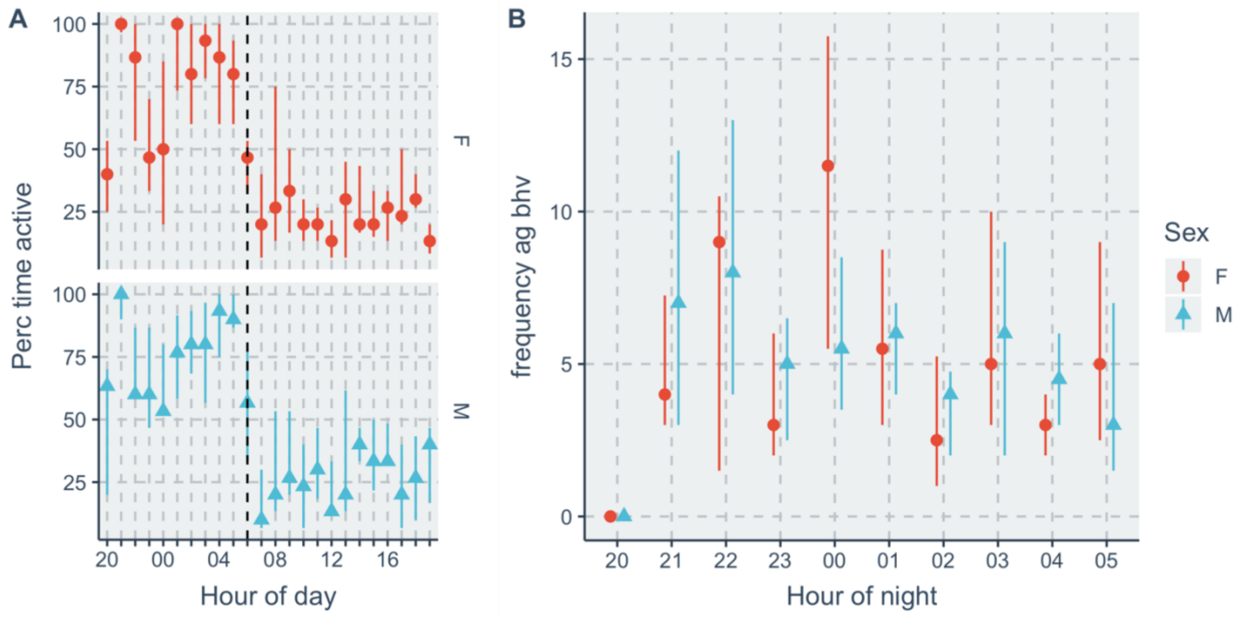


*SI Fig. 3*: General incidence of activity and social behavior. **A**: Scatterplot of median percent time active for each hour of the day for all animals in Batch 1 from the start of darkness (20:00h) to the end of light (19:00h), the vertical-colored lines denote the inter-quartile range, a dashed vertical line denotes time the lights turned on. **B**: Scatterplot of the median frequency of agonistic behavior throughout the night for females and males, the vertical-colored lines denote the inter-quartile range.

*Table 4*: Gene primer sequences

SI Table 7: Gene primer sequences

| **Gene** | **Forward Primer Sequence** | **Reverse Primer Sequence** |
|---|---|---|
| ActinB | CGCCACTAGTTCGACATGGA | AGGGTCAGGATGCCTCTCTT |
| StAR | CCAGCAGGAGAATGGGGATG | TCCTTGACATTTGGGTTCCACT |
| Cyp11A1 | GGCCCCATTTACAGGGAGAA | TACTGGTGATAGGCGACCCA |
| Nrb01 | TGCTCTTTAACCCAGACCTGC | CGGATCTGAGCTGGTACTCTC |
| Nr5a1 | CGAGAGCTGCAAGGGTTTCT | ACCGTCAGGCACTTCTGGA |
| Cyp11b1 | TCTGTGATGTTGCCAGAGGA | AGCCAGGGTTCCAAGATTGT |
